# Elimination of glutamatergic transmission from Hb9 interneurons does not impact treadmill locomotion

**DOI:** 10.1101/2021.01.21.427548

**Authors:** Lina M. Koronfel, Kevin C. Kanning, Angelita Alcos, Christopher E. Henderson, Robert M. Brownstone

## Abstract

The spinal cord contains neural circuits that can produce the rhythm and pattern of locomotor activity. It has previously been postulated that a rhythmogenic population of glutamatergic neurons, termed Hb9 interneurons, contributes to this rhythmogenesis. The homeobox gene, Hb9, is expressed in these interneurons as well as motor neurons. We developed a mouse line in which cre recombinase activity is inducible in neurons expressing Hb9. We then used this line to eliminate vesicular glutamate transporter 2 from Hb9 interneurons, and found that there were no deficits in treadmill locomotion. We conclude that glutamatergic neurotransmission by Hb9 interneurons is not required for locomotor rhythmogenesis. The role of these neurons in neural circuits remains elusive.

## INTRODUCTION

Over the past decade, genetic knowledge has been increasingly harnessed to dissect neural circuits that produce behaviour^1^. This knowledge, for the most part, has been derived from our comprehension of neural development^2^. In the spinal cord, expression patterns of transcription factors have been discovered and cardinal classes of spinal interneurons defined, enabling the development of tools that have been used first to identify and then to functionally alter neuronal populations. These tools have led to concepts of spinal motor circuit organisation and the roles of specific classes of interneurons in producing motor output^3^.

One critical homeobox gene expressed during development is Hb9 (Mnx1 – motor neuron and pancreas homeobox 1). Hb9 is crucial for consolidation of spinal motor neuron (MN) fate during development, and is expressed in somatic MNs as well as spinal visceral MNs (sympathetic preganglionic neurons, SPNs)^4^. Thus a transgenic mouse in which expression of enhanced green fluorescent protein (eGFP) was driven by the promoter for Hb9 (Hb9::eGFP) was made for the selective study of motor neurons^5^. In addition to MNs, a small population of eGFP-expressing interneurons (INs) was seen throughout much of the spinal cord in these transgenic mice^6,7^. With the demonstration that these INs did indeed express endogenous Hb9, they were termed Hb9 interneurons (Hb9 INs)^7^. Thus, it was found that distinct populations of spinal neurons express Hb9: MNs (somatic and SPNs) and Hb9 INs.

As Hb9 INs were shown to be glutamatergic, to be positioned in the ventromedial upper lumbar spinal cord where locomotor rhythm generation occurs^8^, and to have membrane properties that could support pacemaker-type activity, it was proposed that they could have a role in locomotor rhythm generation [^6,7^ reviewed in ^9^]. To test this, it should be possible to study locomotor activity following genetic removal of Hb9 INs from spinal circuits, for example using a binary strategy to eliminate vGluT2 in Hb9-expressing neurons (as was done, for example, in dI3 INs^10^). In fact, it has recently been suggested that doing so leads to impairments in locomotor rhythm^11^.

But there are several problems and potential pitfalls with such an approach. Firstly, it was noted that in Hb9::eGFP transgenic mice, there is GFP expression beyond Hb9 INs and MNs; that is, eGFP expression did not represent “true” Hb9 expression. Furthermore, Cre expression in Hb9^cre^ mice is not limited to Hb9 INs and MNs^11^. Ergo, excision of vGluT2 using Hb9^cre^ mice would not be limited to these neuronal populations, making interpretation of results problematic.

To address this concern, we generated a novel inducible Cre mouse line, Hb9::CreER^T2^, using a BAC transgene. Here, we first characterise the pattern of tamoxifen inducible recombination and demonstrate the specificity and sensitivity of recombination in Hb9-expressing MNs and Hb9 INs. Next, we cross this line with a vGluT2^fl/fl^ line^10,12^ to eliminate vGluT2 expression in Hb9-expressing neurons (and call the line Hb9-vGluT2^OFF^). We then demonstrate that glutamatergic transmission by Hb9 INs does not contribute to treadmill locomotion of varying speeds. We conclude that the role, if any, of Hb9 INs thus remains obscure.

## RESULTS

### Recombination in Hb9::CreER mice is restricted to Hb9-expressing cells

Like others^11^, we initially used an Hb9^cre^ mouse line, but in early experience, found widespread expression of Cre-reporter throughout the embryo (Fig 1A), all levels of the spinal cord, and in neurons throughout all laminae in the spinal cord (Fig 1B; noted over many mice, but n=2 formally analyzed for this study). It was thus clear that we should abandon this strategy.

**Figure 1:**
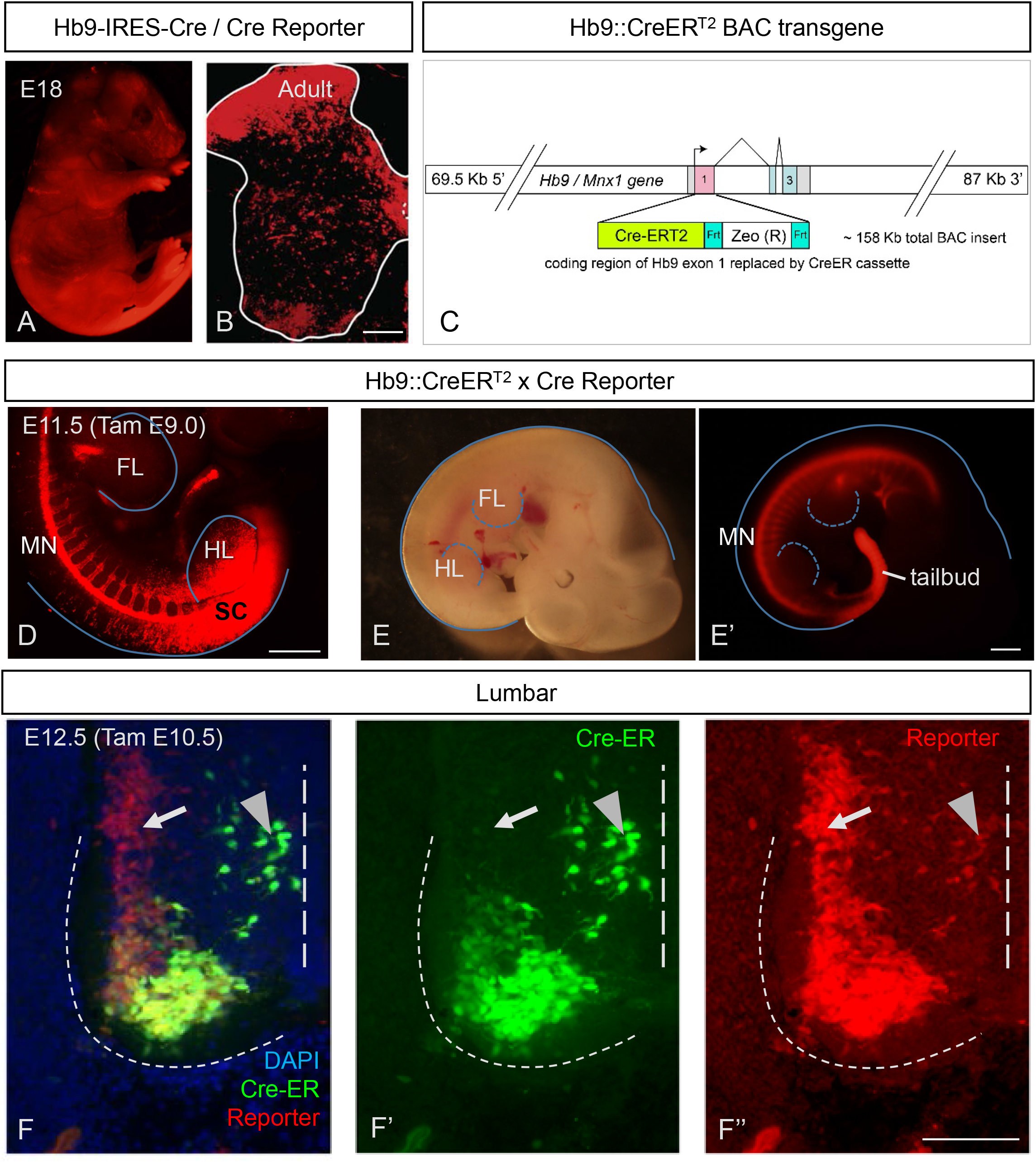
Temporal dependence of Hb9 driven CreER activity on specificity of recombination in embryonic spinal cord. **A-B**: Extensive recombination of Hb9-Cre knock-in mice in many regions, shown in (A) E18 fetus, and (B) transverse postnatal spinal cord. **C**: Diagram of the BAC transgene used to drive CreERT2 expression in Hb9^ON^ cells. **D–F**: Transgenic HB9-CreER^T2^ mice efficiently and specifically enable recombination of Ai14 (tdTom) reporter in HB9^ON^ cells in the embryo, including spinal motor neurons. **D**: Tamoxifen at E9.0 fails to confer MN specific recombination at hindlimb (HL) levels in the spinal cord (SC). **E**: Tamoxifen at E9.5 no longer results in broad SC recombination at hindlimb levels and has activity restricted to the MN column. **F**: Early activation by tamoxifen at E10.5 labels the MN that by E12.5 have downregulated Hb9 (arrow), as reflected by antibody staining for Cre to identify current sites of Cre-ER expression. Arrowhead indicates nascent MN that are still migrating from the midline, but may also include Hb9 INs. Scale bars: B and F, 100 µm; D-E, 500 µm.

We therefore developed an inducible Hb9::CreER^T2^ mouse line (Fig 1C; hereafter called Hb9::CreER), and proceeded to study the selectivity of recombination following administration of tamoxifen (TAM) initially between E9 and E14 in Hb9::CreER;Rosa26-lox-stop-lox-tdTomato (together Hb9::CreER;tdTom). Expression of tdTom was evident as early as 24 hours following TAM. At early embryonic stages (E9; Fig 1D), TAM led to widespread expression of tdTom in the spinal cord, presumably due to early expression of Hb9 outside of the MN lineage in the caudal neural plate at this time point (Fig 1D). This result is similar to the pattern of expression seen in Hb9^cre^;tdTom mice (cf. Figure 1A).

On the other hand, administration of TAM even 12 hours later (E9.5) led to more restricted expression patterns, with clear specificity of recombination (Fig 1E). In particular, ventral horn neurons in the region of motor pools expressed the reporter along with various other neurons (Fig 1F). But it was evident that there were two clusters of tdTom^ON^ MNs, one which maintained Cre-ER expression at E12.5, and one which had very low Cre-ER protein levels (Fig 1F, arrow). These clusters corresponded to two distinct motor columns, the lateral and medial portions of the lateral motor columns (LMC-L and LMC-M), respectively. The populations of somatic MNs that showed recombination dependent expression of tdTom depended on the timing of TAM. While the lateral aspect of the lateral motor column (LMC-L) expressed tdTom with administration any time after E9.5, the medial portion (LMC-M) only expressed tdTom if TAM was given prior to E11.5. This downregulation of Hb9::CreER is consistent with previous observations that endogenous Hb9 expression is down-regulated in LMC-M motor neurons by E12.5 ^13,14^. Note also that motor neurons that have not yet settled in lamina IX express CreER at this embryonic stage (Fig 1F, arrowhead).

Thus, embryonic administration of TAM in Hb9::CreER mice leads to specific activation of Cre recombinase, accurately marking Hb9-expressing motor neurons (Hb9^ON^ MN) in time and space.

Tamoxifen administered at E12 - when MNs are specifically labeled within the spinal cord - also led to reporter expression outside the central nervous system (Fig 2 A-C). Delaying TAM administration until E14 did not alter this non-neuronal recombination (Fig 2 D-I). Sites of recombination included forelimb and hindlimb mesenchyme (Fig 2 A, B, D’), which may correspond to Hb9 expression in developing cartilage (Fig 2D’, E’); gastrointestinal lumen (Fig 2C, I); digit tendons (Fig 2E); notochord (Fig 2F); stomach (Fig2G); and pancreas (Fig 2H). Of note, there was no evidence of tdTom expression in sensory neurons: dorsal root ganglia had no reporter expression.

**Figure 2:**
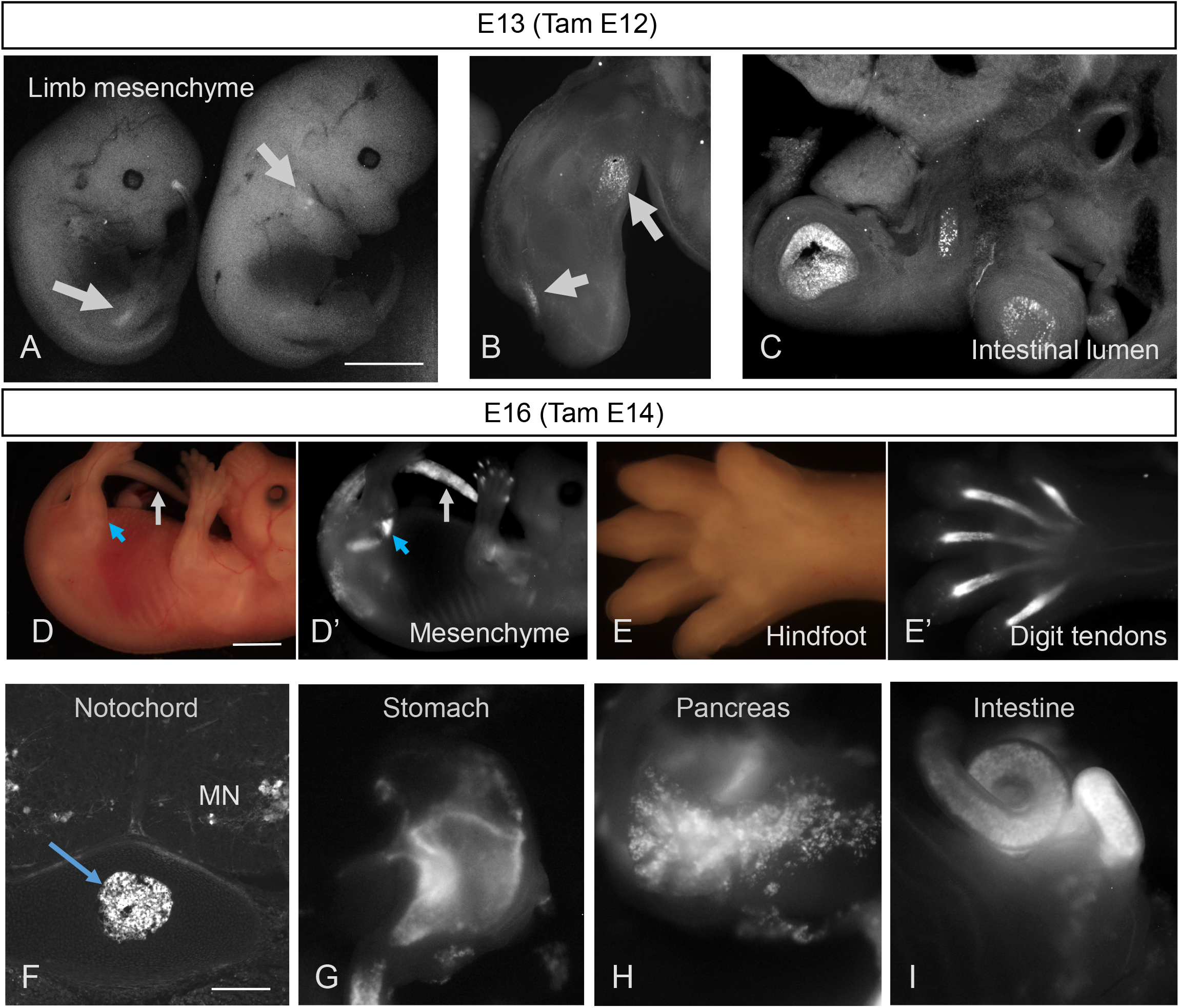
Sites of pre-natal Hb9-CreER recombination outside of the CNS. **A-C**: ROSA-lox-stop-lox-EYFP reporter expression in mid-gestation HB9-CreER mice following IP administration of tamoxifen to pregnant dam at E12, revealing recombination outside of the CNS evident in limbs of E13 animals (A), vibratome sections of forelimb (B), and the lumen of the intestine (C). **D-E**: Brightfield (D, E) versus Ai14 tdTomato reporter expression (D’, E’) in E16 mice induced with tamoxifen at E14, showing recombination in limb and tailbud mesenchyme (D’) and tendons of the knee (D’, blue arrow) and Hindfoot (E’). **F-I**: Additional sites of Ai14 tdTomato reporter expression after E14 tamoxifen induction, including known sites of non-neuronal Hb9 expression such as notochord (F), stomach (G), pancreas (H), and intestine (I). Scale bars: A and D, 2 mm; F, 250 µm.

We next turned to postnatal (up to P7) TAM, which also resulted in efficient recombination in Hb9^ON^ motor neurons: reporter expression in brain stem MNs was specific to Hb9-expressing ventral MNs (vMNs), including abducens (Fig3A) and hypoglossal (Fig3B) nuclei. We noted no other supraspinal expression of reporter except in the choroid plexus. (Note, for example, lack of expression in the facial nucleus (VII), which is of branchial origin, Fig 3A). In spinal MNs, postnatal tamoxifen induced highly efficient recombination in Hb9^ON^ MNs (Fig 3C). During development, LMC-M motor neurons down-regulate Hb9 (see above), and correspondingly these MNs, identified by their location and expression of cholinergic markers VAChT (Fig 3D) or ChAT (Fig 3E), were not labelled with tdTom. However, in Hb9-expressing populations such as the medial motor column (MMC) and LMC-L motor neurons, recombination was efficient, with almost all motor neurons undergoing recombination. In addition, sympathetic preganglionic neurons (SPNs) express Hb9 ^4^, and they too showed recombination following TAM (Fig 3H). (Note that, consistent with patterns of Hb9 expression, other cholinergic neurons did not.)

**Figure 3:**
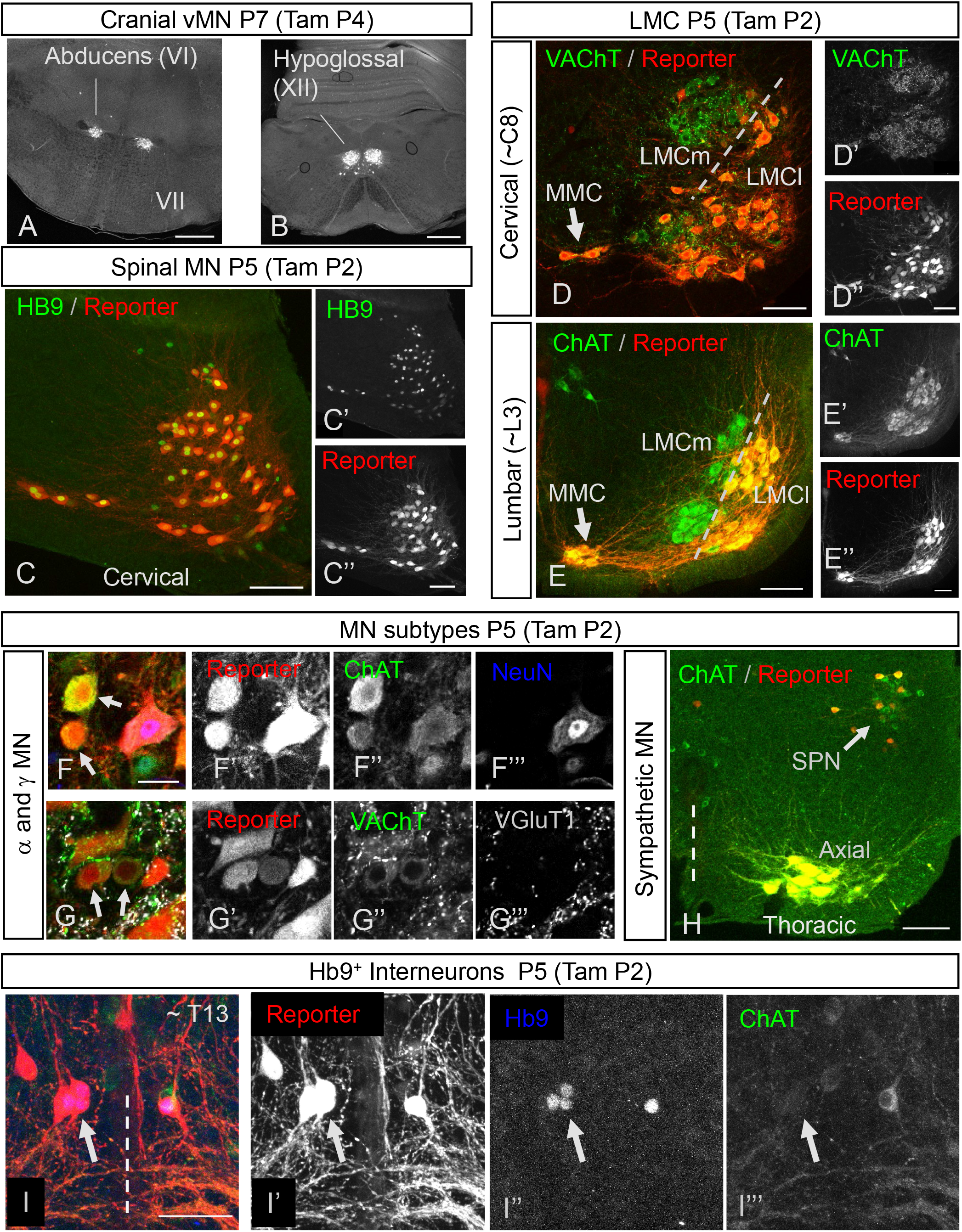
Postnatal tamoxifen induced recombination specific to Hb9^ON^ neurons. **A-B**: Ai6 Cre-reporter expression in vMN cranial nuclei following P4 Tamoxifen administration to Hb9::CreER mice, showing specific recombination in motor nuclei VI (A) and XII (B) but no other brainstem MN such as VII (A). **C**: In spinal cord, tamoxifen administration at P2 results in high correspondence between recombination in Ai14 reporter (C, C’) and endogenous Hb9 protein expression. **D-E**: Consistent with Hb9 expression patterns, post-natal Hb9::CreER recombination distinguishes LMC medial and lateral sub-divisions at both cervical (D) and lumbar (E) levels, as revealed by Ai14 reporter pattern in all MN visualized with either VAChT (D) or ChAT (E), following tamoxifen induction at P2. **F-G**: Both α- and γ-MNs undergo efficient recombination in Hb9-CreER mice following tamoxifen administration at P2. α-MN are identified as larger MN that are NeuN^ON^ (F), or with dense VGluT1 and VAChT puncta on their soma (G). γ-MN are smaller MN that are NeuN^OFF^ (arrows in F; scale bar= 25 µm), or with little to no VGluT1 and VAChT puncta on their soma. **H**: Sympathetic Preganglionic Neurons (SPNs) at thoracic levels of spinal cord undergo recombination in Hb9::CreER mice following tamoxifen administration at P2. **I**: Hb9 INs in Hb9::CreER mice efficiently undergo inducible recombination following tamoxifen at P2 (Level T13 ~ L1 shown). Hb9 INs (arrow) are identified by size and position near the central canal, expression of Hb9 protein (I, I’’) and lack of ChAT. Scale bars: A-B, 500 µm; C,D,E, and H, 100 µm; F (also applies to G) and I, 50 µm.

We next examined Hb9::CreER post-natal recombination in α- and γ-MN subtypes, as non-BAC based Hb9::GFP transgenic mice display selective expression in α-MN^15^. We identified γ-MNs as small, ChAT^ON^ neurons with a visible nucleus that did not express NeuN (Fig 3F), or those that had a paucity of primary afferent (labelled by vGluT1) or C-bouton (labelled by vAChT) inputs (Fig 3G). Using postnatal TAM regimens, we identified efficient recombination in both NeuN^OFF^, VGluT1^low^, VAChT^low^ γ-MNs as well as NeuN^ON^, VGluT1^high^, VAChT^high^ α-MNs (Fig3 F, G). Therefore, within Hb9^ON^ motor columns the Hb9::CreER BAC line efficiently recombines in all MN subtypes.

We next turned our attention to Hb9 interneurons: a population of glutamatergic neurons in medial lamina VIII above the second lumbar segment (L2)^7^. Following postnatal TAM administration, there was CreER activation in this region as noted by reporter expression (Fig 3I). These neurons were positive for Hb9 and negative for ChAT, indicating that these are Hb9 INs (Fig 3I). Virtually all visually-identified Hb9 INs, identified by their location, size, and Hb9 expression, expressed the reporter (see β-gal expression below for quantification).

Together, these data reflect the specificity of our approach, in which recombination is seen primarily during the times when, and in the neurons in which, endogenous Hb9 is expressed. That is, recombination in Hb9::CreER mice induced by TAM any time after E10.5 and including the early post-natal period was seen in MNs that endogenously express Hb9: LMC-L MNs, MMC MNs, SPNs, vMN brain stem MNs, and Hb9 INs. But when TAM was administered prior to E11.5, there was also recombination in LMC-M MNs (Fig 1F). Administration prior to E9.5 led to widespread, non-specific recombination in caudal regions (Fig 1D). These data indicate that these inducible Hb9::CreER mice can be used for spatially and temporally specific and sensitive recombination strategies aimed at Hb9-expressing MNs and Hb9 INs.

### vGlut2-dependent transmission by Hb9 INs does not affect treadmill locomotion

As our goal was to study mature mice, we next ensured that expression induced by postnatal TAM administration led to persistent and specific expression in adult mice (Fig 4A). When crossed into Hb9^lacZ^ knock-in mice^16^, reporter expression in the adult was essentially identical to β-gal expression (205/216 tdTom neurons in 3 mice also expressed β-gal), and was evident in MNs, SPNs, and Hb9 INs (Fig 4A,B). We did not find β-gal-expressing neurons that did not express tdTom. That is, recombination was efficient at this age, and there was no evidence of spurious off-target recombination.

**Figure 4:**
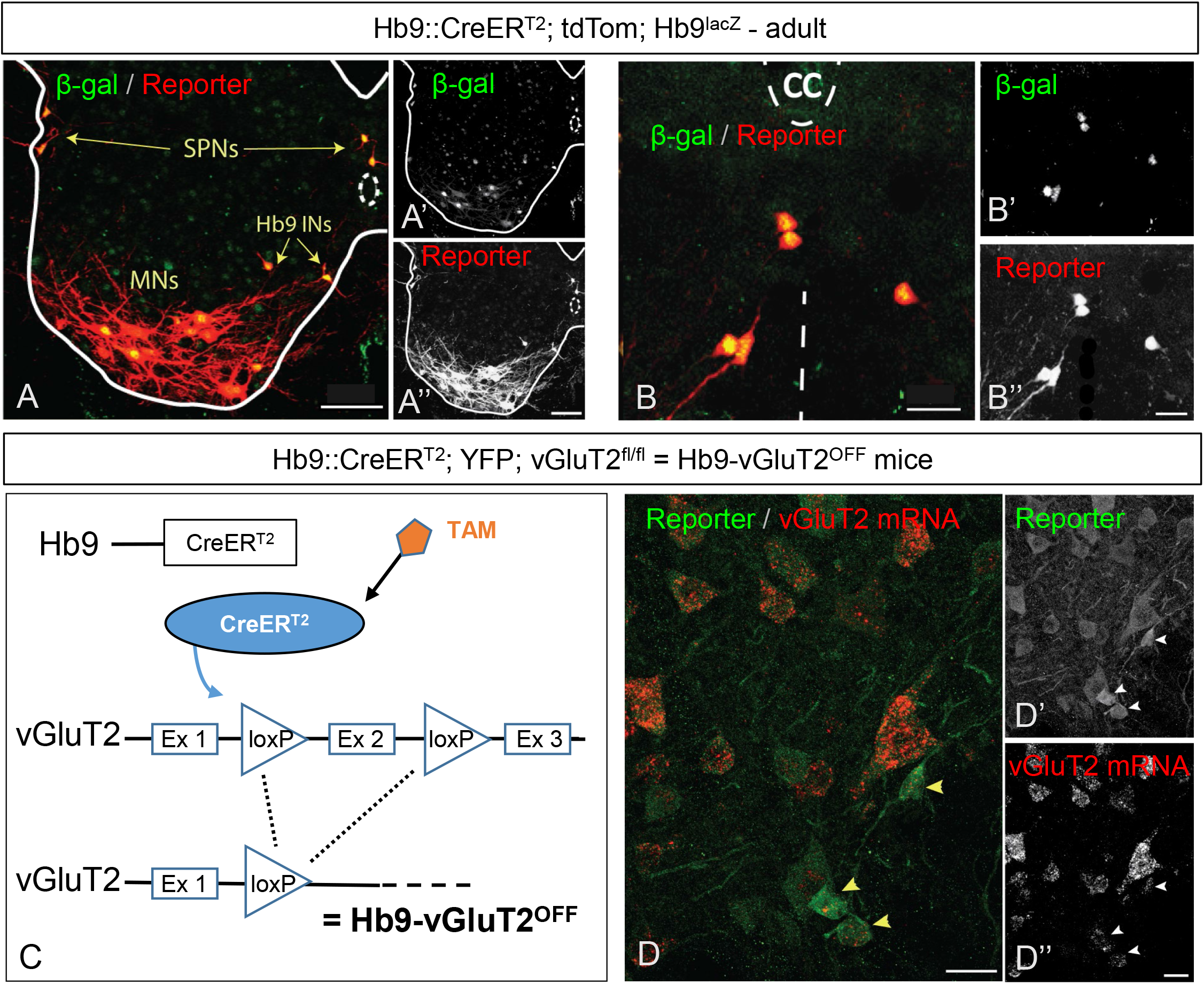
Conditional recombination in adult Hb9::CreER mice. **A**. Following tamoxifen administration to Hb9^lacZ^;Hb9::CreER;R26-lox-stop-lox-tdTom adult mice, there is overlap of expression of β-gal and td-Tom reporter in specific neurons: SPNs, MNs, and Hb9 INs. **B**. Higher magnification image of Hb9 INs in the ventromedial region of upper lumbar spinal segment. **C**. Illustration of conditional CreER-mediated excision of vGluT2 in Hb9 INs following tamoxifen administration, creating Hb9-vGluT2^OFF^ mice (TAM: tamoxifen, Ex: exon). **D**. Fluorescent in situ hybridization showing elimination of vGluT2 mRNA in Hb9::CreER; R26-lox-stop-lox-YFP; vGluT2^fl/fl^ mice. Hb9 INs (arrowheads) lack vGluT2 mRNA following recombination. Scale bar: A, 50 µm; B and D, 20 µm.

Given that Hb9 interneurons are glutamatergic ^7^, whereas the primary neurotransmitter of MNs is acetylcholine, we reasoned that we could functionally remove Hb9 INs from circuits by crossing Hb9::CreER mice with vGluT2^flox/flox^ mice^12^, to yield Hb9-vGluT2^OFF^ mice (Fig 4C), in which glutamatergic transmission from neurons expressing Hb9 would be eliminated ^10^. In situ hybridisation for vGluT2 mRNA showed that this strategy was effective (n=2; Fig 4D; cf. Fig 3A in ^7^).

We next assessed these mice in the fourth and sixth post-natal weeks at various treadmill locomotor speeds, quantifying a number of locomotor parameters as well as their variability (Fig 5). Parameters related to the pattern of locomotion were the same in control (n=7, of which n=5 could maintain the highest speed) and Hb9-vGLuT2^OFF^ (n=11, of which n=9 could maintain the highest speed) mice. That is, at all speeds at which the mice could consistently locomote, the means of fore-hind limb (Fig 5A top) and right-left (Fig 5B top) coupling, and rear track width (Fig 5C top) were all the same in Hb9-vGluT2^OFF^ and control mice (Table 1). Furthermore, the variability in these parameters was identical between the two groups of mice (fore-hind limb coupling, right-left coupling, and rear track width in lower panels of Fig 5A-C, respectively). These data indicate that glutamatergic transmission by Hb9 INs does not contribute to the generation of the locomotor pattern required for treadmill locomotion.

**Table 1:**
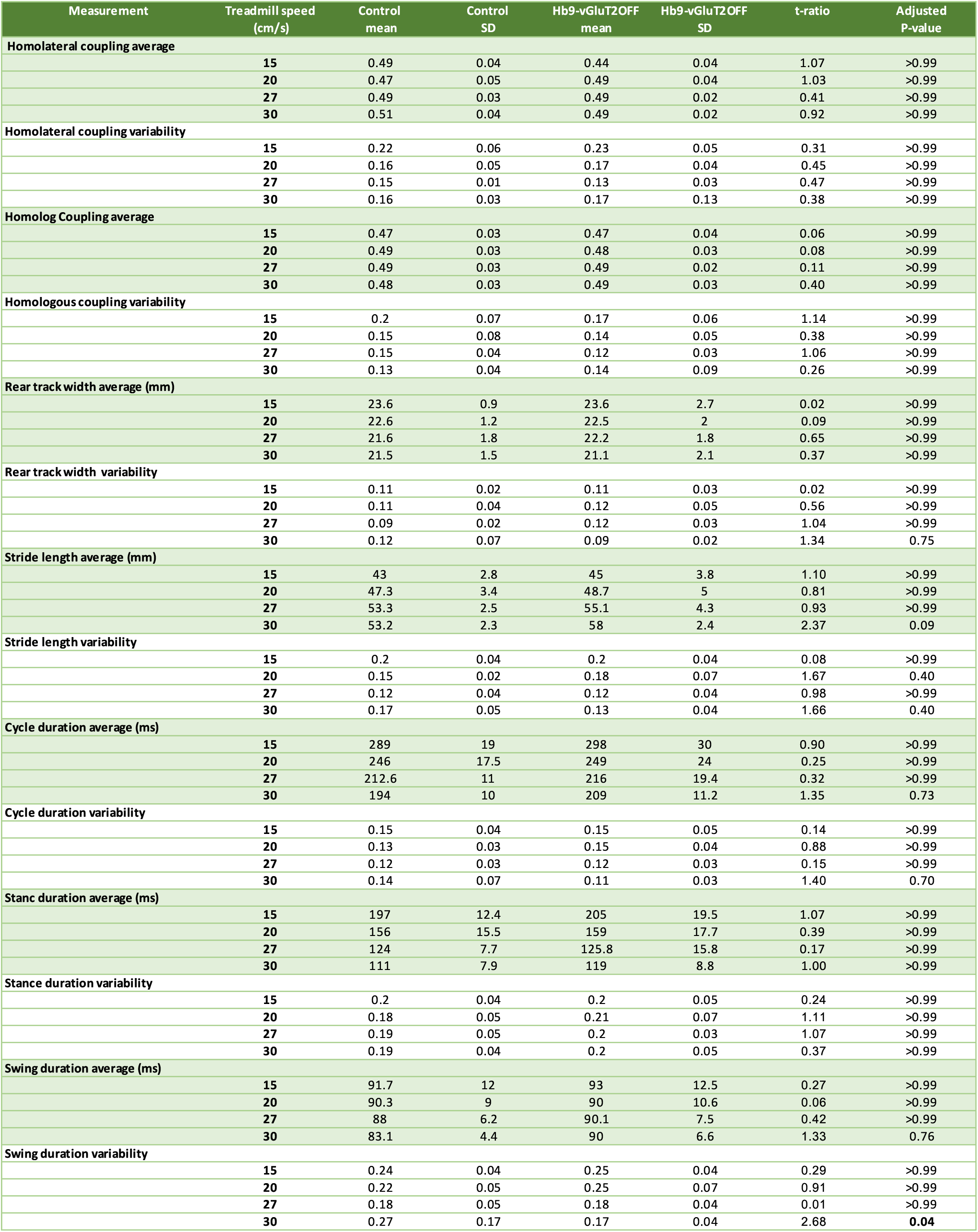
Statistics table for all treadmill parameters. Bonferroni-Dunn corrected p-values. Degrees of Freedom: 56. See Methods for details.

**Figure 5:**
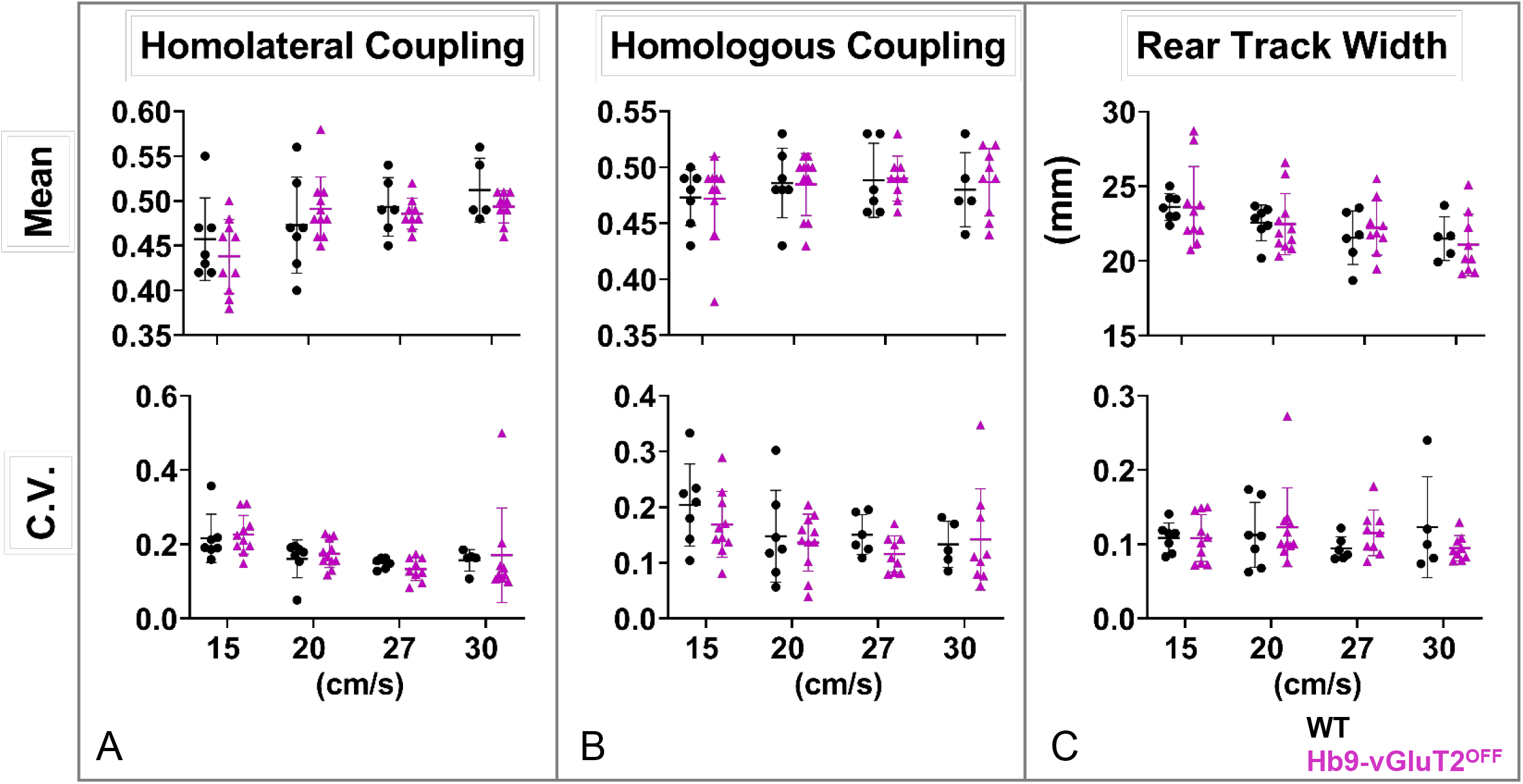
The pattern of locomotion in Hb9-vGluT2^OFF^ mice is similar to that of controls. **A**. No significant difference was observed in homolateral coupling compared to WT when comparing the means (upper panel, P>0.99 at all speeds examined). There were no differences in the coefficients of variation either (lower panel, P>0.99 at all speeds). **B**. Similarly, neither homologous coupling (upper panel) nor its coefficients of variation (lower panel, P>0.99 at all speeds) were different. **C**. Rear track widths were also no different (upper panel, P>0.99 for all speeds), with similar coefficients of variation (lower panel, P>0.99 for all speeds except 30 cm/s: P=0.75). Numbers of mice used in these experiments were total of 7 control mice: n= 7, 7, 6, and 5 for 15, 20, 27, and 30 cm/s, respectively, and total of 11 Hb9-vGluT2^OFF^ mice: n= 10, 11, 9, and 9 for the 4 speeds, respectively. Details of statistics are in Table 1.

As we and others ^6,7,9,11,17,18^ had previously suggested that Hb9 INs may provide a “clock” function for locomotor circuits, we suspected there may be differences, in particular increased variability, in timing parameters in Hb9-vGluT2^OFF^ mice. We could not detect any significant differences (Table 1) in stride length (Fig 6A top), step cycle duration (Fig 6B top), or extensor/flexor timing, with stance (Fig 6C top) and swing (Fig 6D top) durations being identical. Also, the relationships between these phases and cycle duration (Fig 7A) were the same in Hb9-vGluT2^OFF^ and control mice. Furthermore, the coefficients of variability for each of these parameters (Figure 6, lower panels) were also the same between the different groups, with the only one in any doubt being the coefficient of variability of the swing phase at the fastest speed (p=0.04). Similarly, when these experiments were repeated in early post-weaned mice (P21-23), no differences were found in the Hb9-vGLuT2^OFF^ mice (Fig 7B), suggesting that the “normality” seen two weeks later was not a result of post-natal compensation. That is, the absence of glutamatergic transmission by Hb9 INs did not influence locomotor rhythmicity.

**Figure 6:**
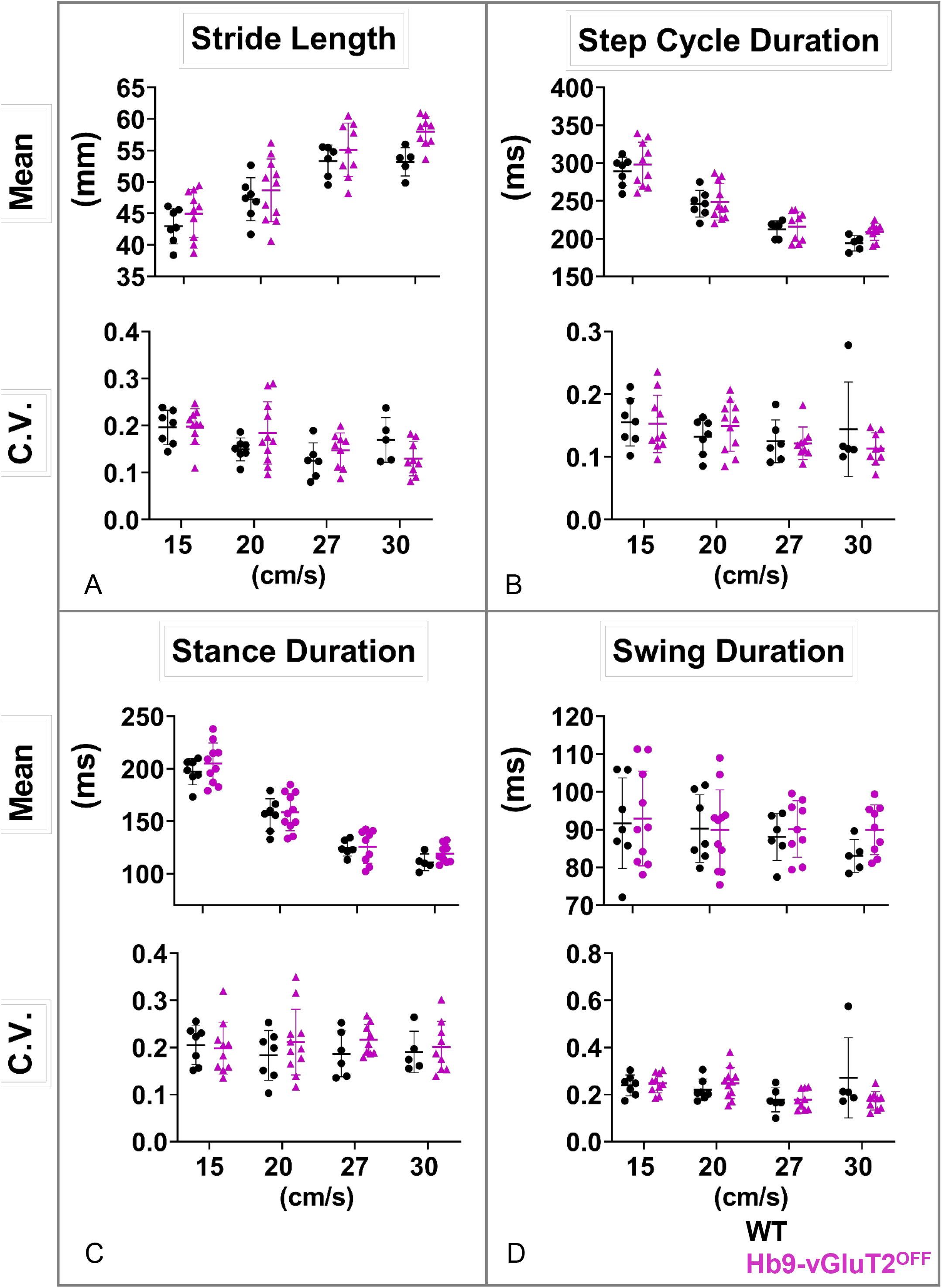
The rhythm of locomotion in Hb9-vGluT2^OFF^ mice is similar to that of controls. **A**. Stride length was the same (upper panel, P>0.99 at all speeds except 30 cm/s, which was P=0.09), as well as coefficients of variation (lower panel, P>0.99 at 15 and 27 cm/s and P=0.4 at 20 and 30 cm/s). **B**. Similarly, no difference was observed in cycle duration (upper panel, P>0.99 at 15, 20, and 27 cm/s and P=0.73 at 30 cm/s). Coefficients of variation, were also the same (lower panel, P>0.99 at 15, 20, and 27 cm/s, and P=0.7 at 30 cm/s). **C**. Stance durations were also similar (upper panel, P>0.99 at all speeds), as well as their coefficients of variation (lower panel, P>0.99 at all speeds). **D**. There was no difference in swing duration (upper panel, P>0.99 at speeds 15, 20, and 27 cm/s, and P=0.76 at 30 cm/s). The coefficients of variation were the same at 15, 20, and 27 cm/s (lower panel, P>0.99), but were significantly lower in Hb9-vGluT2^OFF^ mice at 30 cm/s (P=0.04), seemingly due to an outlier in the control group. Number of mice used at the 4 speeds for the control group (total 7) are 7, 7, 6, and 5, and for the Hb9-vGluT2^OFF^ group (total 11) are 10, 11, 9, and 9, respectively. Details of statistics are in Table 1.

**Figure 7:**
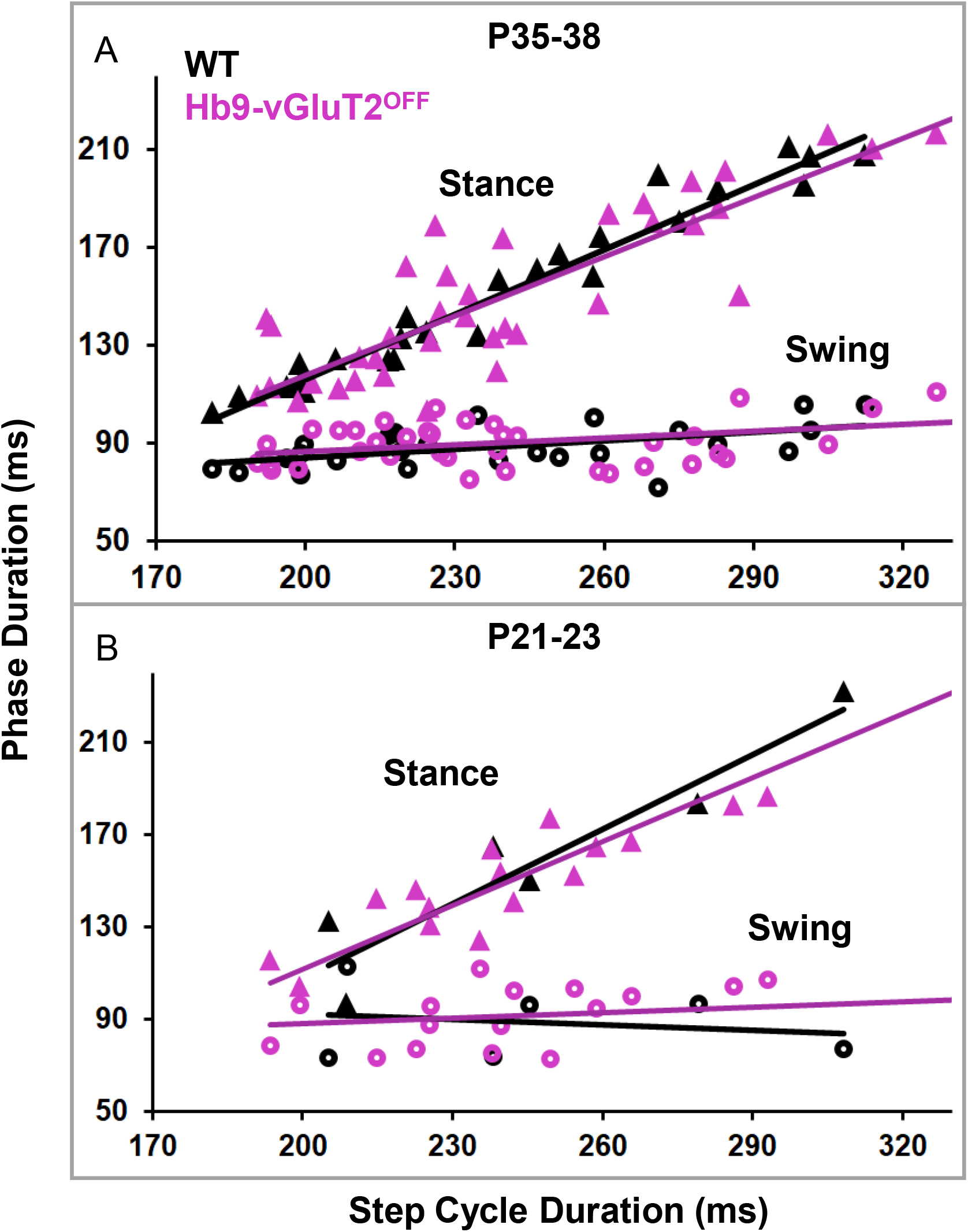
Relationships between phase duration and step cycle duration in Hb9-vGluT2^OFF^ mice were similar to controls. **A**. In P35-38 mice, there was no significant difference in the relationship between stance phase and speed (WT R^2^=0.96, Hb9-vGluT2^OFF^ R^2^=0.79, P=0.42), and no difference in swing phase in relation to speed (WT R^2^=0.27, Hb9-vGluT2^OFF^ R^2^=0.16 P=0.65). For the following treadmill speeds 15, 20, 27, and 30 cm/s, WT n=7, 7, 6, and 5, respectively (total n=7), whereas for Hb9-vGluT2^OFF^ n=10, 11, 9, and 9, respectively (total n=11). **B**. Similarly, In P20-23 mice, there were no significant differences in stance phase (WT R^2^=0.88, Hb9-vGluT2^OFF^ R^2^=0.96, P=0.32) or swing phase (WT R^2^=0.04, Hb9-vGluT2^OFF^ R^2^=0.14, P=0.32) in relation to speed. For the following treadmill speeds 10, 15, and 20 cm/s, WT n=2, 2, and 2 respectively (total n=2). Whereas for Hb9-vGluT2^OFF^ n=8, 8, and 7, respectively (total n=8).

Taken together, we were unable to find evidence to support the hypothesis that glutamatergic transmission by Hb9 INs plays a significant role in treadmill locomotion.

## DISCUSSION

Hb9 is a homeodomain protein expressed in sub-populations of somatic motor neurons and a population of ventral spinal interneurons termed Hb9 interneurons whose function is unknown. In order to gain genetic access to these neurons, we made a new BAC transgenic mouse line using a tamoxifen-inducible Cre recombinase strategy, Hb9::CreER. After demonstrating the specificity of this line, we proceeded to ask whether Hb9 INs significantly contribute to locomotor activity in adult mice. To do this, we removed glutamatergic neurotransmission from Hb9 INs using an intersectional approach to delete vGluT2 from Hb9-expressing neurons. We demonstrated that doing so does not significantly impact treadmill locomotion.

### Use of Hb9::CreER mice to study motor neurons

Several methods have been used to gain genetic access to somatic motor neurons, but all – including ours – have caveats. The mouse line that we describe here can be used to study and manipulate LMC-L MNs, or with precise timing of TAM administration at E9, possibly even all MNs. While the limitation to LMC-L would be disadvantageous for many studies, it could provide the ability to compare affected (LMC-L) vs unaffected (LMC-M) motor neurons in the same animal with any given genetic manipulation. This could be useful, for example, in ALS studies^19^.

In contrast to our line, Hb9-ires-Cre knock-in mice (Mnx1^tm4(cre)Tmj^, termed Hb9^cre^ here) have been used in several studies to genetically manipulate cervical motor neurons ^20^;^21^. To ensure specificity, these investigators used an intersectional approach. Specifically, Hb9-cre;Isl2-lox-stop-lox-DTA (or DTX) crosses were used to eliminate most cervical motor neurons ^21^;^20^. Of note, the focus in those studies was the neuromuscular junction. This intersectional approach would not affect neuronal populations other than motor neurons as isl2-expressing INs do not express Hb9^21^. Hb9-driven Cre expression was not extensively investigated in these studies, but appeared to be confined to lamina IX (motor neurons) in a section of cervical spinal cord; this may have led others to the erroneous assumption that Hb9-ires-Cre was motor neuron specific throughout the spinal cord (see http://www.informatics.jax.org/reference/allele/MGI:2447793?typeFilter=Literature).

Several years later, Cre-mediated recombination was studied in these Hb9^cre^ mice, and a gradient of non-MN recombination was evident: there was reporter expression in relatively few cells outside motor pools in the cervical cord, but extensive expression throughout the lumbar cord (^22^;^23,24^. That is, there is expression of Cre beyond MN pools in these mice, and this “ectopic” expression increases from rostral to caudal. This pattern likely results from early Hb9 expression in the caudal neural plate and tail bud area, where all cell progeny derived from these neural plate progenitor cells inherit Cre-recombined DNA. Moreover, this line shows expression in supraspinal regions. Although other genetic approaches enable Cre-mediated recombination in motor neurons (Olig2^cre^ or ChAT^cre^), both Olig2 and ChAT are expressed in other populations of spinal (ChAT) and supraspinal (both) neurons (e.g.,^25^). Further, neither can be induced in the post-natal period (TAM needs to be given embryonically in Olig2-CreER mice, and in ChAT-CreER mice, expression is not confined to motor neurons^26^. Thus, the Hb9::CreER line provides improved genetic access to non-LMC-M spinal motor neurons, as well as other Hb9-expressing neurons including SPNs and Hb9 INs.

### Locomotor rhythmogenesis

Our primary goal in this study was to test our long-standing hypothesis that Hb9 INs are involved in locomotor rhythmogenesis. Hb9 INs were identified almost 2 decades ago, and their rhythmic, conditional bursting properties along with their location in the ventromedial upper lumbar spinal cord^3^ led to the suggestion that they perform a “clock” role to maintain the rhythm of locomotion^7,9^. Since that time, these neurons have been studied in a number of additional labs as well ^17,18,27^, but their function in behavior has remained opaque.

Using the Hb9^cre^ knock-in line, it was suggested that eliminating glutamatergic transmission from Hb9 expressing neurons impacted locomotor rhythm in isolated neonatal spinal cord preparations^11^. But as they and we showed, it is difficult to draw conclusions from this approach, as early expression of Hb9 leads to Cre-mediated recombination in many neurons through the spinal cord, i.e. Cre recombination is too widespread to draw solid conclusions (see above). Thus, to allow more specific genetic access, it was necessary first to make a new mouse that could obviate the problems resulting from early caudal Hb9 expression.

Using an inducible CreER system, we indeed found specific and sensitive recombination in Hb9 expressing neurons when activated with appropriate timing of TAM administration.

With this new mouse, we generated Hb9-vGluT2^OFF^ mice by multi-generational breeding with vGluT2^flox^ mice, which we and others have used successfully before ^3,10,12^, and used these mice to quantify treadmill locomotion in the absence of glutamatergic transmission from Hb9 INs. The pattern and rhythm of treadmill walking was virtually identical in mutant and control mice. Of all parameters studied, the only statistical difference seen was a decrease in the variability of swing duration, which resulted from a single outlier mouse in the control group.

It is important to note, however, that these data do not completely exclude Hb9 INs from participating in locomotion. For example, our protocol involved post-natal TAM administration, and Hb9 INs could be important during embryonic development. More interestingly, however, Hb9 INs are electrotonically coupled to other types of neurons^16^; electrotonic transmission can play an important role in synchronicity and rhythmogenesis in other systems (e.g. ^28^;^29^;^30^). The Hb9::CreER mouse described here could be used to try to sort this out, for example by post-natally eliminating Cx36 from Hb9-expressing neurons, when motor neuron expression of Cx36 is already decreasing^31^. Selectively eliminating Hb9 INs using, for example, genetic expression of diphtheria toxin fragment A^32^ driven by the promoter for vGluT2 could also be used.

If Hb9 INs are not responsible for locomotor rhythmogenesis, then which neurons fulfill this role? Hb9 INs seemed like the perfect candidate neurons to maintain the rhythm of locomotion. Other candidate neurons, including Shox2 INs, have been proposed as playing this role^30^. And while Hb9 INs may still contribute to rhythmogenesis via electrotonic coupling, it is interesting to consider parallels with respiratory rhythmogenesis. There have been discussions over decades about whether the respiratory rhythm is generated by pacemaker neurons and/or circuit mechanisms, with no single neuronal substrate found^33–35^. Perhaps the locomotor field will face these same debates. Nonetheless, it is interesting to consider that two populations of neurons that reciprocally inhibit each other are a prime substrate for rhythm generation^36^, and that locomotor circuits have many places for such pairs of direct or indirect reciprocal inhibition (governing movement across joints, between joints, and between limbs, for example^37^. Thus there are many microcircuits in which rhythm could be generated with or without neuronal pacemaker properties. In retrospect, perhaps, it would have been a surprise to see profound changes in rhythm when removing chemical transmission from this one neuron type – Hb9 INs – from spinal cord circuits.

## Methods

### Animals

All experimental procedures at Dalhousie University were approved by the University Committee on Laboratory Animals and were in accordance with the Canadian Council on Animal Care guidelines and those procedures at Columbia University were approved in accordance with IACUC guidelines.

Hb9^lacZ/+ 4^, Hb9^cre/+ 4^, and vGluT2^flox/flox 12^ have been previously described. Rosa26-lox-stop-lox-reporters (tdTomato (Ai14, Jax#007908); EYFP (Jax#006148); or ZsGreen (Ai6; Jax#007906)) were obtained from the Jackson Laboratory.

### Hb9::CreER^T2^ DNA Recombineering

The bacterial artificial chromosome (BAC) clone RP24-351I23 (CHORI BACPAC resources) containing the mouse *Mnx1* (Hb9) gene was modified using recombineering bacterial strains (NCI, http://web.ncifcrf.gov/research/brb/recombineeringInformation.aspx). The BAC was modified to remove the loxP and loxP511 sites in the pTARBAC vector backbone using homologous targeting cassettes containing ampicillin and spectinomycin resistance, respectively. A shuttle vector containing short homology arms to desired recombination sites was used to introduce CreER^T2^-BGH polyA and a Frt-Zeo-Frt selection cassette into Hb9 exon 1, replacing the Hb9 coding region. In the modified BAC, the CreER^T2^ cassette^38^ was flanked by 69.5 kilobases of 5’ DNA and 87 kilobases of 3’ DNA. The Frt-Zeo-Frt cassette is retained in the transgene but lacks a eukaryotic promoter.

Both Dalhousie University and UCL have material transfer agreements with both Columbia University, where the mice were engineered, and Novartis Forschungsstiftung, Zweigniederlassung, Friedrich Miescher Institute for Biomedical Research (“FMI”), for the CreER^T2^.

### Transgenic mouse generation

Modified BAC DNA was digested by Not1 to remove pTARBAC vector, run out on low melting point agarose, the insert gel purified, and recovered by beta-agarase digestion. DNA was dialyzed into injection buffer and oocyte injections were done in the transgenic core at the Irving Cancer Center, Columbia University Medical Center. Three Hb9::CreER founder lines (4, 8, and 10) were confirmed for tamoxifen induced recombination in motor neurons between E11 – E14 using a ROSA26-lox-stop-lox-EYFP reporter. Line 10 gave low level inducible activity and was terminated. Line 4 and Line 8 displayed similar inducible recombination activity in the desired cell populations and were maintained. Both lines are viable if homozygosed, but were maintained as heterozygotes. Because the lines were indistinguishable, we have used line 8 for the majority of studies described herein.

### Tamoxifen Administration

Tamoxifen (Sigma T-5648) was dissolved to 20 mg/mL in 90% sesame oil/ 10% Ethanol, briefly heated to 50°C to facilitate dissolution, and stored in frozen aliquots. For administration, stock aliquots were warmed to 37°C for 10 min prior to injection. Pregnancies were timed by vaginal plugs, and pregnant dams (~30g bodyweight) were intraperitoneally (IP) injected a single time with 0.1 mL stock (2 mg) at indicated ages. In our preliminary characterization of the mice using the ROSA26::lox-stop-lox-EYFP reporter, we found that a single intraperitoneal injection administered between E10 and E16 of tamoxifen into a pregnant female Hb9::CreER mouse was highly efficient at inducing recombination in embryonic spinal motor neurons.

In postnatal experiments, pups of both sexes (ages P2 – P7) were injected subcutaneously above the dorsal neck fat pad with 0.1mL (2mg) stock solutions and maintained on a 37°C heating pad for 1 hour before returning to their nursing cage. Due to leakage from the subcutaneous injection site, the exact dose in pups was likely variable. We did not observe any toxicity when tamoxifen was administered by these approaches.

### Tissue preparation and labeling

Under deep ketamine/xylazine anesthesia, mice were perfused with 4% paraformaldehyde (PFA) in 0.2M PB. Following perfusion, the spinal cords were extracted and post fixed in PFA at 4oC overnight, then transferred to a 30 % sucrose solution for 24-48 hours before sectioning.

Spinal cord blocks were sectioned (50-70 μm) using a vibrating microtome and processed for immunohistochemistry on the same day, or stored in glycerol at −20° C for later use.

Floating sections were washed in PBS for 10 minutes before incubation in 50% ethanol for 30 minutes to enhance antibody (Ab) penetration. Sections were washed with double salt PBS (dsPBS) 3 × 10 minutes and transferred to blocking solution containing 10% donkey serum in 0.3M Triton X and PBS solution (PBST) for 30 minutes at room temperature.

Tissues were transferred to primary Ab (Anti-DsRed-Rabbit, Rockland antibodies and assays PA, USA 1:2000; Anti-β-gal Goat or Mouse, Santa Cruz Biotechnology, Inc, Dallas, Tx, USA 1:500; Anti-GFP-Sheep or Mouse, GenWay Biotech, Inc. San Diego, CA, USA and Novus Biological inc. Littleton, CO, USA 1:500) diluted in a solution containing 1% donkey serum and PBST and incubated for 48-72 hours at 4o C. After incubation, sections were washed in dsPBS 3 × 10 minutes and transferred to secondary Ab solution to be incubated overnight at 4oC. Secondary Abs (Alexa 546 Donkey anti-Rabbit; Alexa 488 Donkey anti-goat, anti-mouse, or anti-sheep, all 1:400, Invitrogen, Oregon, USA) were diluted in a solution containing 1% donkey serum and PBST. Finally, sections were washed in PBS for 10 minutes and mounted using Vectashield mounting medium (Vector laboratories, Burlingame, CA).

### In situ hybridization combined with immunohistochemistry

Tissues were processed for combined fluorescent *in situ* hybridization and immunohistochemistry as previously described^7,10^, and based on^39^. The primer for antisense digoxigenin (DIG) riboprobe for vGluT2 was provided by the Jessell lab (Columbia University, USA), and the probe was kindly prepared by the Fawcett lab (Dalhousie University, NS, Canada). Briefly, sections were fixed in 4% PFA for 10 minutes at RT followed by washing in PBS 3 × 3 minutes. Sections were acetylated for 10 minutes in an acetic anhydride buffer and transferred to a hybridization solution (50% formamide, 5xSSC, 5xDenhardt’s, 250 µg/mL Baker’s yeast RNA, 500 µg/mL salmon sperm DNA; Sigma), initially without the probe at RT overnight. Sections were then heated in diluted hybridization solution with the probe for 5 minutes at 80°C, then quickly transferred to ice before being incubated at 72°C overnight in the solution. On day three, sections were incubated in 0.2x saline sodium citrate buffer (SSC) at 72°C 2 × 30 minutes followed by equilibrating the sections in 0.2x SSC for 5 minutes at RT. Sections were incubated in tris NaCl blocking buffer (0.1M Tris chloride, 0.15M NaCl, 0.5% blocking reagent; TNB) for 1 hour at RT, then overnight in a solution containing donkey serum, sheep anti-DIG-POD (1:500 TNB; Roche), and rabbit anti-GFP antibody (1:500 TNB; Chemicon) at 4°C. On day four, tyramide signal amplification (with AlexaFluor 555 or Cy3 (Perkin-Elmer), TSA kit #42, Invitrogen) was used to boost the fluorescent in situ signal. Sections were then incubated in anti-rabbit-Alexa 488 (Invitrogen) antibody for 3 hours in PBS and mounted using Vectashield mounting medium (Vector laboratories, Burlingame, CA).

### Image acquisition

Confocal images were acquired with either a Zeiss LSM 710 Laser Scanning Confocal Microscope (Dalhousie) or Leica TCS SP5 (Columbia). For imaging synaptic puncta (VGlut1, VAChT), vibratome sections were imaged in Z stacks with a 20x or 63X objective (2µm optical Z resolution). Images were captured either as 2D snapshots or 3D z-stacks with intervals of 0.8-1.88 μm and total Z-thickness up to approximately 30 μm. Pinhole size was set to 1 airy unit for all channels.

### Treadmill locomotion

Locomotor behaviour was studied on a treadmill (Cleversys, Inc.) equipped with a transparent belt and a high speed camera (Basler, USA). Mice were placed in a 161 cm^2^ chamber, and gait was recorded with the supplied software (BCam Capture Version 2.00, Cleversys, Inc.) at frame rate of 100 frames/s. The treadmill belt and chamber were cleaned with Peroxigard prior to every experimental session. Locomotor behaviour was examined at belt speeds of 10, 15, and 20 cm/s for the age group of postnatal day 21-23 (7.7 ± 1.9g), and speeds of 15, 20, 27, and 30 cm/s for age groups of postnatal day 35-38 (15.9 ± 1.7g). Faster speeds were attempted, however mice could not maintain locomotion at those speeds (e.g. 40 cm/s and 50 cm/s). The speeds were chosen to represent low, medium, and high speeds respectively^40^. Video recording started as soon as the treadmill speed was stable for 20 seconds per trial and mice were allowed to rest for 1-2 minutes between each trial. In some cases, they were studied on a second day to confirm observations.

### Gait analysis

Gait parameters were analyzed using TreadScan Version 3.00 (Cleversys, Inc.). Prior to each experiment, a ruler was placed along the treadmill and imaged for scale calibration. An artificial foot model was created by drawing a polygon over the foot of interest at different frames, and saving the RGB ratios that represent each foot. To exclude the instances when the mouse did not keep up with speed (e.g. when rearing or grooming), we included only sequences during which the mouse was keeping up with the treadmill, and thus was at the front of the treadmill chamber. For each step, analysis parameters were obtained including: stance time, swing time, cycle duration, stride length, homologous coupling, homolateral coupling, and rear track width. Stride length, defined as the distance travelled by the limb in a complete step cycle, was calculated as (running sped * stride time) + displacement, where displacement could be positive or negative depending on the change of position of the foot in the camera frame. The reliability of each step was confirmed by examining all frames to ensure that each foot was adequately captured before exporting the raw data to an Excel file for analysis. The right hindlimb was used to quantify single limb parameters, both hindlimbs for homologous, and right fore- and hind-limbs for homolateral coupling.

### Statistics

For each step cycle parameter (Figure 5 and 6), unpaired 2-tailed t-tests were used (based on the assumption that samples at each speed were obtained from populations with the same scatter, as borne out by CV analysis), followed by Bonferroni-Dunn post-hoc correction for multiple comparisons. For the comparison of swing or stance vs cycle duration (Figure 7), linear regression analysis was done using ANCOVA-type calculations to compare ^41^. All tests were done using Prism (GraphPad, San Diego, CA 8.4.3(686)). P-values < 0.05 were considered to be significant.

## Acknowledgments

We would like to thank Boris Lamotte d’Incamps for early and helpful electrophysiological experiments with these mice, Tom Jessell’s lab for the generous gifts of Hb9^lacZ^ and Hb9^cre^ mice as well as the in situ probe, Jim Fawcett’s lab for preparing the in situ probe, and Nadia Farbstein for technical support. We also thank Victor Lin in the Columbia Transgenic core for technical guidance in preparing the BAC transgene, the Motor Neuron Center at Columbia University for equipment use, and Pierre Chambon for CreER^T2^ plasmid. This work was supported by grants to RMB from the Canadian Institutes of Health Research (MOP 79413) and Wellcome Trust (110193). RMB’s position is supported by Brain Research UK. KCK and CEH received support from NIH/NINDS grants F32 NS055547 and 1R01NS056422, with additional support to CEH from the Claire and Leonard Tow Foundation and the SMA Foundation.

## Author Contributions

KCK and CEH made the BAC transgene and established the Hb9::CreER mouse. KCK did experiments and analysed data for Figures 1–3. LMK collected mouse behavioural data, analysed it, and made Figures 4–7 except for in situ experiments, which were done and analysed by AA. LMK, KCK, and RMB conceived the experiments and drafted the article, which was critically revised and then approved by all authors.

## Additional Information

The authors have no competing interests to declare. All work done by KCK and CEH was while at Columbia University.

## Notes

### Competing Interest Statement

The authors have declared no competing interest.

## Reference List

1 Grillner, S. & Jessell, T. M. Measured motion: searching for simplicity in spinal locomotor networks. Curr Opin Neurobiol 19, 572–586, doi:10.1016/j.conb.2009.10.011 (2009).

2 Alaynick, W. A., Jessell, T. M. & Pfaff, S. L. SnapShot: spinal cord development. Cell 146, 178-178.e171 (2011).

3 Kiehn, O. Development and functional organization of spinal locomotor circuits. Current Opinion in Neurobiology 21, 100–109, doi: https://doi.org/10.1016/j.conb.2010.09.004 (2011).

4 Arber, S. et al. Requirement for the homeobox gene Hb9 in the consolidation of motor neuron identity. Neuron 23, 659–674 (1999).

5 Wichterle, H., Lieberam, I., Porter, J. A. & Jessell, T. M. Directed differentiation of embryonic stem cells into motor neurons. Cell 110, 385–397 (2002).

6 Hinckley, C. A., Hartley, R., Wu, L., Todd, A. & Ziskind-Conhaim, L. Locomotor-like rhythms in a genetically distinct cluster of interneurons in the mammalian spinal cord. Journal of neurophysiology 93, 1439–1449 (2005).

7 Wilson, J. M. et al. Conditional rhythmicity of ventral spinal interneurons defined by expression of the Hb9 homeodomain protein. J Neurosci 25, 5710–5719 (2005).

8 Kjaerulff, O. & Kiehn, O. Distribution of networks generating and coordinating locomotor activity in the neonatal rat spinal cord in vitro: a lesion study. Journal of Neuroscience 16, 5777–5794 (1996).

9 Brownstone, R. M. & Wilson, J. M. Strategies for delineating spinal locomotor rhythm-generating networks and the possible role of Hb9 interneurones in rhythmogenesis. Brain Res Rev 57, 64–76 (2008).

10 Bui, T. et al. Circuits for grasping: spinal dI3 interneurons mediate cutaneous control of motor behavior. Neuron 78, 191–204, doi:10.1016/j.neuron.2013.02.007 (2013).

11 Caldeira, V., Dougherty, K. J., Borgius, L. & Kiehn, O. Spinal Hb9::Cre-derived excitatory interneurons contribute to rhythm generation in the mouse. Scientific Reports 7, 41369, doi:10.1038/srep41369 (2017).

12 Hnasko, T. S. et al. Vesicular glutamate transport promotes dopamine storage and glutamate corelease in vivo. Neuron 65, 643–656 (2010).

13 Rousso, D. L., Gaber, Z. B., Wellik, D., Morrisey, E. E. & Novitch, B. G. Coordinated actions of the forkhead protein Foxp1 and Hox proteins in the columnar organization of spinal motor neurons. Neuron 59, 226–240 (2008).

14 Dasen, J. S., De Camilli, A., Wang, B., Tucker, P. W. & Jessell, T. M. Hox repertoires for motor neuron diversity and connectivity gated by a single accessory factor, FoxP1. Cell 134, 304–316 (2008).

15 Shneider, N. A., Brown, M. N., Smith, C. A., Pickel, J. & Alvarez, F. J. Gamma motor neurons express distinct genetic markers at birth and require muscle spindle-derived GDNF for postnatal survival. Neural development 4, 1–22 (2009).

16 Wilson, J. M., Cowan, A. I. & Brownstone, R. M. Heterogeneous Electrotonic Coupling and Synchronization of Rhythmic Bursting Activity in Mouse Hb9 Interneurons. Journal of Neurophysiology 98, 2370–2381, doi:10.1152/jn.00338.2007 (2007).

17 Tazerart, S., Vinay, L. & Brocard, F. The persistent sodium current generates pacemaker activities in the central pattern generator for locomotion and regulates the locomotor rhythm. J Neurosci 28, 8577–8589 (2008).

18 Kwan, A., Dietz, S., Webb, W. & Harris-Warrick, R. Activity of Hb9 interneurons during fictive locomotion in mouse spinal cord. The Journal of neuroscience: the official journal of the Society for Neuroscience 29, 11601–11613, doi:10.1523/jneurosci.1612-09.2009 (2009).

19 Orr, B. O. et al. Presynaptic Homeostasis Opposes Disease Progression in Mouse Models of ALS-Like Degeneration: Evidence for Homeostatic Neuroprotection. Neuron 107, 95–111 e116, doi:10.1016/j.neuron.2020.04.009 (2020).

20 Pun, S. et al. An intrinsic distinction in neuromuscular junction assembly and maintenance in different skeletal muscles. Neuron 34, 357–370 (2002).

21 Yang, X. et al. Patterning of muscle acetylcholine receptor gene expression in the absence of motor innervation. Neuron 30, 399–410, doi:10.1016/s0896-6273(01)00287-2 (2001).

22 Kramer, E. R. et al. Cooperation between GDNF/Ret and ephrinA/EphA4 signals for motor-axon pathway selection in the limb. Neuron 50, 35–47, doi:10.1016/j.neuron.2006.02.020 (2006).

23 Li, X.-M. et al. Retrograde regulation of motoneuron differentiation by muscle β-catenin. Nature neuroscience 11, 262–268 (2008).

24 Hess, D. M. et al. Localization of TrkC to Schwann cells and effects of neurotrophin-3 signaling at neuromuscular synapses. J Comp Neurol 501, 465–482 (2007).

25 Ju, J. et al. Olig2 regulates Purkinje cell generation in the early developing mouse cerebellum. Sci Rep 6, 30711, doi:10.1038/srep30711 (2016).

26 Rotolo, T., Smallwood, P. M., Williams, J. & Nathans, J. Genetically-directed, cell type-specific sparse labeling for the analysis of neuronal morphology. PLoS One 3, e4099 (2008).

27 Anderson, T. et al. Low-threshold calcium currents contribute to locomotor-like activity in neonatal mice. Journal of Neurophysiology 107, 103–113, doi:10.1152/jn.00583.2011 (2012).

28 Eisen, J. S. & Marder, E. A mechanism for production of phase shifts in a pattern generator. J Neurophysiol 51, 1375–1393 (1984).

29 Rekling, J. C., Shao, X. M. & Feldman, J. L. Electrical coupling and excitatory synaptic transmission between rhythmogenic respiratory neurons in the preBotzinger complex. J Neurosci 20, RC113 (2000).

30 Ha, N. T. & Dougherty, K. J. Spinal Shox2 interneuron interconnectivity related to function and development. Elife 7, e42519 (2018).

31 Chang, Q. & Balice-Gordon, R. J. Gap junctional communication among developing and injured motor neurons. Brain Res Brain Res Rev 32, 242–249, doi:10.1016/s0165-0173(99)00085-5 (2000).

32 Ivanova, A. et al. In vivo genetic ablation by Cre-mediated expression of diphtheria toxin fragment A. Genesis 43, 129–135, doi:10.1002/gene.20162 (2005).

33 Anderson, T. M. & Ramirez, J.-M. Respiratory rhythm generation: triple oscillator hypothesis. F1000Research 6 (2017).

34 Feldman, J. L. & Kam, K. Facing the challenge of mammalian neural microcircuits: taking a few breaths may help. The Journal of physiology 593, 3–23 (2015).

35 Ramirez, J. M. & Baertsch, N. Defining the Rhythmogenic Elements of Mammalian Breathing. Physiology (Bethesda) 33, 302–316, doi:10.1152/physiol.00025.2018 (2018).

36 Kristan, W. B. & Katz, P. Form and function in systems neuroscience. Curr Biol 16, R828–831 (2006).

37 Danner, S. M., Shevtsova, N. A., Frigon, A. & Rybak, I. A. Computational modeling of spinal circuits controlling limb coordination and gaits in quadrupeds. Elife 6, doi:10.7554/eLife.31050 (2017).

38 Feil, R., Wagner, J., Metzger, D. & Chambon, P. Regulation of Cre Recombinase Activity by Mutated Estrogen Receptor Ligand-Binding Domains. Biochemical and Biophysical Research Communications 237, 752–757, doi:https://doi.org/10.1006/bbrc.1997.7124 (1997).

39 Vosshall, L. B., Amrein, H., Morozov, P. S., Rzhetsky, A. & Axel, R. A spatial map of olfactory receptor expression in the Drosophila antenna. Cell 96, 725–736, doi:10.1016/s0092-8674(00)80582-6 (1999).

40 Beare, J. E. et al. Gait analysis in normal and spinal contused mice using the TreadScan system. J Neurotrauma 26, 2045–2056, doi:10.1089/neu.2009.0914 (2009).

41 Zar, J. H. Biostatistical analysis. (Prentice-Hall, 1974).

